# Shared PKS modules in biosynthesis of synergistic laxaphycins

**DOI:** 10.1101/2020.05.29.098152

**Authors:** LMP Heinilä, DP Fewer, J Jokela, M Wahlsten, A Jortikka, K Sivonen

## Abstract

Cyanobacteria produce a wide range of lipopeptides that exhibit potent membrane-disrupting activities. Laxaphycins consist of two families of structurally distinct macrocyclic lipopeptides that act in a synergistic manner to produce antifungal and antiproliferative activities. Laxaphycins are produced by range of cyanobacteria but their biosynthetic origins remain unclear. Here, we identified the biosynthetic pathways responsible for the biosynthesis of the laxaphycins produced by *Scytonema hofmannii* PCC 7110. We show that these laxaphycins, called scytocyclamides, are produced by this cyanobacterium and are encoded in a single biosynthetic gene cluster with shared polyketide synthase enzymes initiating two distinct non-ribosomal peptide synthetase pathways. To our knowledge, laxaphycins are the first clearly distinct polyketide synthase and non-ribosomal peptide synthetase hybrid natural products with shared branched biosynthesis. The unusual mechanism of shared enzymes synthesizing two distinct types of products may aid future research in identifying and expressing natural product biosynthetic pathways and in expanding the known biosynthetic logic of this important family of natural products.

## Introduction

Natural products are chemical compounds produced by living organisms, with research interest focused on discovery of new natural products with pharmaceutical applications (Spainhour, 2005; Newman and Cragg, 2016). Many bioactive natural products have complex chemical structures with rare chemical moieties that allow them to react with specific molecular targets and to kill or inhibit the growth of other organisms (Rodrigues et al., 2016). Cyanobacteria produce a wide variety of natural products with different activities and complicated structures (Demay et al., 2019; Huang and Zimba, 2019). Characterization of new natural products offers starting material for drug design as new active structures (Rodrigues et al., 2016). Characterization of the biosynthesis of these products advance methods in production of the structures through combinatorial biosynthesis (Kim et al., 2015). Many microbial and cyanobacterial natural products are constructed by polyketide synthases (PKS) and non-ribosomal peptide synthetases (NRPS) (Kehr et al., 2011; Dittmann et al., 2015). PKS and NRPS enzymes often act together and are encoded in joint gene clusters producing hybrid PKS/NRPS products (Miyanaga et al., 2018). PKS and NRPS enzymes allow the production of complex structures with characteristic non-proteinogenic amino acids and the combination of non-ribosomal peptides (NRP) with polyketide chains and decorations (Evans et al., 2011). NRPS and PKS biosynthesis typically follow a co-linearity rule, where the genomic order of the catalytic domains corresponds to the order of the product structure (Guenzi et al., 1998; Callahan et al., 2009). The natural product family of laxaphycins are hypothesized to be produced by the PKS/NRPS hybrid pathway (Bornancin et al., 2015; Bornancin et al., 2019).

Laxaphycins are cyanobacterial cyclic lipopeptides that fall in two distinct structural macrocycles consisting of either 11 amino acids (known as A-type laxaphycins) or 12 amino acids (known as B-type laxaphycins). Both types include β-aminooctanoic acid (Aoa) or β-aminodecanoic acid (Ada)(Table 1). Eleven- and 12-residue laxaphycins have strong synergistic activity in antifungal and antiproliferative bioactivity assays (Frankmölle et al., 1992b; MacMillan et al., 2002; Cai et al., 2018). The biosynthetic origins of members of the laxaphycin family remains unclear. Despite sharing the same name, they are chemically distinct and are anticipated to be produced by distinct pathways. The nomenclature of laxaphycins is complicated due to the two distinct core types addressed as a single family. Furthermore, naming new members after the producing organisms and distinguishing variants with lettering is a poor indication of which type the variant belongs to. We refer to the two types as 11- and 12-residue laxaphycins. There are 30 diverse members assigned to the laxaphycin family reported to date (Table 1). The first laxaphycins to exhibit synergistic effects were described from *Anabaena laxa* (Frankmölle et al., 1992a; Frankmölle et al., 1992b). Here, we focused on laxaphycin variants called scytocyclamides produced by *Scytonema hofmannii* PCC 7110. *S. hofmannii* PCC 7110 was previously studied by our group and a methanol crude extract of the cells was antifungal but the active agent was not identified (Shishido et al., 2015).

**Table 1.**
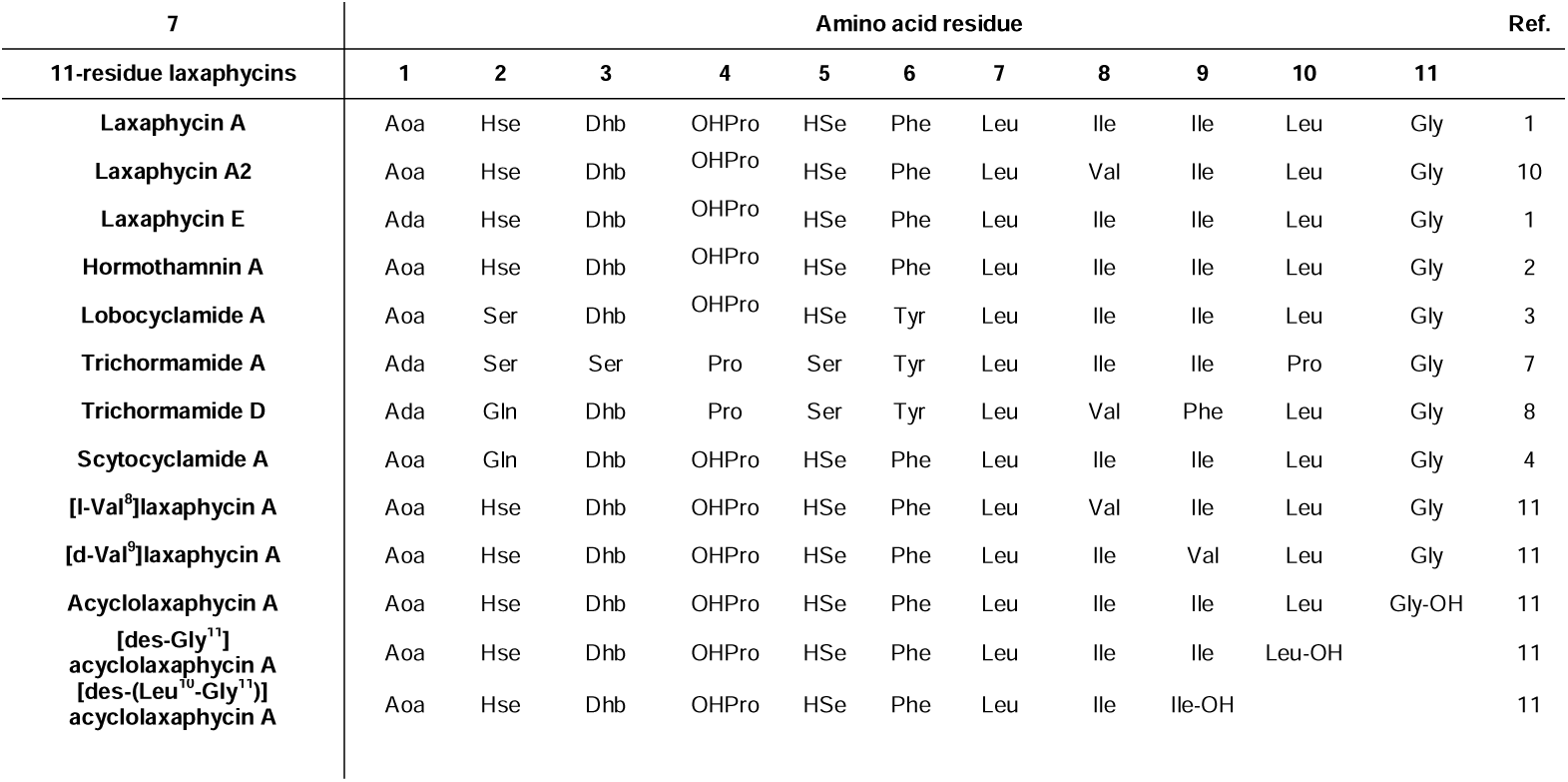
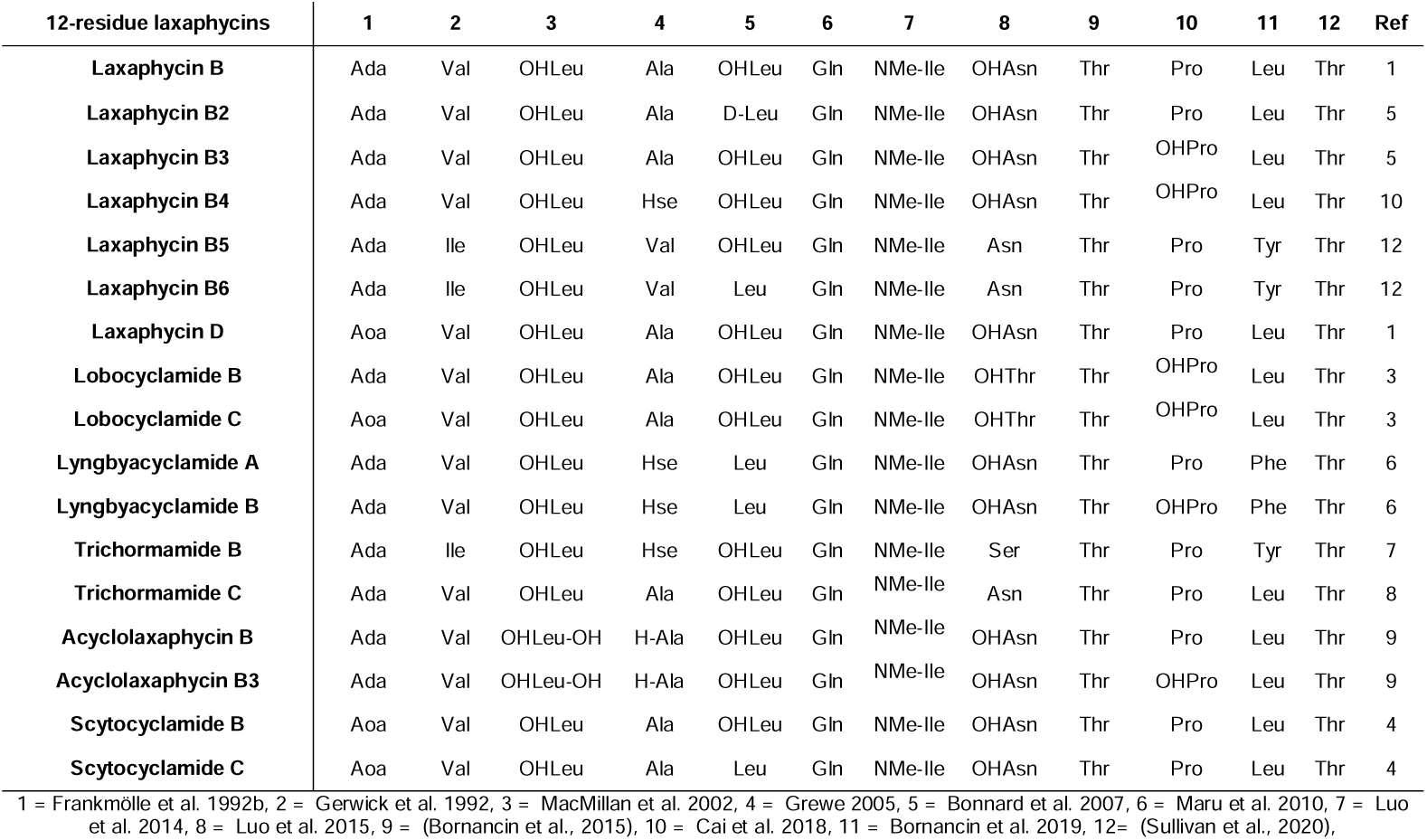
Amino acid sequence of laxaphycin variants.

In this study, we describe the biosynthetic pathways responsible for the biosynthesis of scytocyclamides from *S. hofmannii* PCC 7110. We show that the two types of scytocyclamides are produced by shared PKS enzymes. These enzymes initiate two distinct NRPS pathways, which are exceptional to the PKS/NRPS colinearity rule. We also report the synergistic antifungal activity of scytocyclamides and three new laxaphycin variants (scytocyclamides A2, B2, and B3).

## Materials and Methods

### Scytocyclamide purification

*S. hofmannii* PCC 7110 was grown in 5-L Erlenmeyer flasks with 2.7 L modified Z8 medium without source of combined nitrogen at 20-21**°**C with photon irradiation of 3-7 μmol(m)-2(s)-1 with constant sterilized air bubbling for 3-5 weeks. Cells were collected by decanting excess media and centrifugation at 8000 × g for 5 min. Cells were frozen at -80**°**C and freeze-dried with CHRIST BETA 2-8 LSC plus with a LYO CUBE 4-8 freeze drier. The total amount of freeze-dried biomass was 4 g.

For each gram of dry cells, 30 ml of methanol was used and the mixture was homogenized with Heidolph Silentcrusher M at 20 000 rpm for 30 s. The suspension was centrifuged 10 000 × g for 5 min and supernatant was collected. Extraction of the precipitate was repeated with 30 ml of methanol. Chomatorex (Fuji-Davison Chemical Ltd., Aichi, Japan) chromatography silica ODS powder (10 ml) was added to the supernatant pool and the mixture was dried with rotary evaporator Büchi Rotavapor R-200 at 30**°**C. Solid phase extraction (SPE) was performed with Phenomenex SPE strata SI-1 silica 5 g/20 ml column, preconditioned with 20 ml isopropanol and 20 ml of heptane. Silica ODS powder with the dry extract was added on top of the column and extracted with heptane, ethyl acetate, acetone, acetonitrile, and methanol with each fraction collected individually. Fractions were dried with nitrogen gas flow and re-dissolved in 1 ml of methanol for bioactivity assays. The active methanol fraction was further fractionated with liquid chromatography. Chromatography was performed with an Agilent 1100 Series liquid chromatograph with a Phenomenex Luna 5 *µ*m C18(2) (150 × 10 mm, 100 Å) column. The sample was injected in 100-*µ*l batches and eluted with acetonitrile/isopropanol 1:1 (solvent B) and 0.1% HCOOH (solvent A) with a flow rate of 3 ml min^−1^ in the following four stages: 1, isocratic stage of 43% solvent B in A for 15 min; 2, a linear gradient of solvent B from 43% to 60% in 10; 3, a linear gradient of solvent B from 60% to 81% in 5 min; and 4, a linear gradient of solvent B from 81% to 100% in 6 min. Six scytocyclamide fractions were collected, dried with nitrogen, and weighed.

### 3-hydroxyleucine feeding experiment

*S. hofmannii* PCC 7110 was grown in 100-mL Erlenmeyer flasks with 41 mL modified Z8 medium without a source of combined nitrogen with 40 *µ*M of racemic 3-hydroxyleucine mixture of all four isomers (2-Amino-3-hydroxy-4-methylpentanoic acid, ABCR) to determine if 3-hydroxyleucine is utilized as a substrate in scytocyclamide production. Control cultivations were grown on the same medium without added 3-hydroxyleucine. For both media, three duplicates were cultivated at 20-21 **°**C with photon irradiation of 3-7 μmol(m)-2(s)-1 for 17 days. Cells were collected by decanting excess media and centrifugation 8000 × g for 5 min. Cells were frozen at -80**°**C and freeze-dried with CHRIST BETA 2-8 LSC plus with a LYO CUBE 4-8 freeze drier. Freeze-dried biomass was weighed and extracted with 0.5 ml methanol and glass beads (0.5-mm glass beads, Scientific Industries Inc, USA) using a FastPrep cell disrupter two times for 25 s at a speed of 6.5 m/s. Samples were centrifuged at room temperature for 5 min at 10 000 × g and supernatant was collected.

### Peptide identification by LC-MS

*S. hofmannii* PCC 7110 was grown in 500-mL Erlenmeyer flasks of with 250 mL modified Z8 medium without a source of combined nitrogen at 20-21**°**C with photon irradiation of 3-7 μmol(m)-2(s)-1 with constant sterilized air bubbling for 4 weeks. Cells were collected by decanting excess media and centrifugation at 8000 × g for 5 min. Cells were frozen at -80**°**C and freeze-dried with CHRIST BETA 2-8 LSC plus with a LYO CUBE 4-8 freeze drier. Freeze-dried cells (100 mg) were extracted with 1 ml methanol and glass beads (0.5-mm glass beads, Scientific Industries Inc, USA) using a FastPrep cell disrupter two times for 25 s at a speed of 6.5 m/s. Samples were centrifuged at room temperature for 5 min at 10 000 × g. Supernatant was collected and extraction was repeated with 1 ml of methanol.

Extracts and purified scytocyclamide methanol solutions were analyzed with UPLC-QTOF (Acquity I-Class UPLC-SynaptG2-Si HR-MS, Waters Corp., Milford, MA, USA) equipped with a KinetexLJ C8 column (2.1 × 50 or 100 mm, 1.7 *µ*m, 100 Å, Phenomenex, Torrance, CA, USA) injected with 0.5 or 1 *µ*l samples, eluted at 40°C with 0.1 % HCOOH in water (solvent A) and acetonitrile/isopropanol (1:1, + 0.1 % HCOOH, solvent B) with a flow rate of 0.3 ml min^−1^. Two solvent gradients were used: 1, 5% B to 100% B in 5 min, maintained for 2 min, back to 5% B in 0.50 min, and maintained for 2.50 min before next run; and 2, 10% B to 70% of B in 5 min, then to 95% of B in 0.01 min, maintained for 1.99 min, then back to 10% of B in 0.5 min, and finally maintained for 2.5 min before the next run. QTOF was calibrated using sodium formate and UltramarkLJ 1621, which yielded a calibrated mass range from *m/z* 91 to 1921. Leucine Enkephalin was used at 10-s intervals as a lock mass reference compound. Mass spectral data were accumulated in positive electrospray ionization resolution mode. The MS^E^ Trap Collision Energy Ramp Started from 40.0 eV and ended at 70.0 eV.

### Bioactivity assays

The same *S. hofmannii* PCC 7110 methanol extract used for LC-MS was used for antimicrobial activity screening. The screening was performed with fungal and bacterial strains reported in Table 2. The following samples were pipetted directly on spots on agar: 50 *µ*l cyanobacterial cellular methanol extract, 50 *µ*l negative control (methanol), and 10 *µ*l positive control (nystatin) (Nystatin, *Streptomyces noursei*, EMD Millipore Corp, Germany) solution 5 mg/ml in methanol for fungi and 10 *µ*l ampicillin (Ampicillin sodium salt, Sigma, Israel) 50 mg/ml in 70% ethanol for bacteria. Solvents were allowed to evaporate, leaving the extracts diffused in the agar. Inocula were prepared by growing the fungi for 2-14 days on PDA (Potato Dextrose Agar) media at 28°C and bacteria for two days on BHI (Brain Heart Infusion) agar at 37°C. Cell mass was transferred with a cotton swab from the agar to 3 ml of sterile 5 M NaCl solution or sterile water in the case of *A. flavus*. The inocula were spread on the agar with cotton swabs. Fungal plates were incubated at 28°C and bacterial plates at 37°C for 2 days and analyzed for inhibition zones.

**Table 2.** Strains used in bioassays

| Organism | Strain | Medium | Incubation temperature |
| --- | --- | --- | --- |
| **Fungi** |  |  |  |
| *Candida albicans* | FBCC 2462 | PDA | 28°C |
| *Candida guilliermondii* | FBCC 2457 | PDA | 28°C |
| *Candida krusei* | FBCC 2464 | PDA | 28°C |
| *Candida parapsilosis* | FBCC 2465 | PDA | 28°C |
| *Filobasidiella neoformans* | FBCC 2466 | PDA | 28°C |
| *Aspergillus niger* | FBCC 2467 | PDA | 28°C |
| *Aspergillus parasiticus* | FBCC 2500 | PDA | 28°C |
| *Aspergillus flavus* | FBCC 2467 | PDA | 28°C |
| **Bacteria** |  |  |  |
| *Staphylococcus aureus* | HAMBI 66 | BHI | 37°C |
| *Enterococcus faecium* | HAMBI 1821 | BHI | 37°C |
| *Bacillus cereus* | HAMBI 1881 | BHI | 37°C |
| *Micrococcus luteus* | HAMBI 2688 | BHI | 37°C |
| *Pseudomonas aeruginosa* | HAMBI 25 | BHI | 37°C |
| *Escherichia coli* | HAMBI 1723 | BHI | 37°C |
| *Actinobacter baumannii* | HAMBI 1760 | BHI | 37°C |
| *Enterobacter aerogenes* | HAMBI 1898 | BHI | 37°C |
| *Salmonella enterica* | HAMBI 2331 | BHI | 37°C |

The antifungal activity of purified scytocyclamide fractions dissolved in methanol were tested with *A. flavus* performed as with the cellular extract. Disc diffusion assays were performed with purified scytocyclamides. Paper discs (Blank monodiscs, Abtek biologicals Ltd, UK) were prepared with methanol solutions of the peptides, methanol as a negative control, and nystatin as a positive control. *A. flavus* inoculum was prepared as previously and spread on the plate. Disks were placed on agar and the plates were incubated at 28°C for 2 days and analyzed.

### Biosynthetic gene cluster analysis

The *S. hofmannii* PCC 7110 draft genome sequence (ANNX02) was analyzed with AntiSMASH 4.1 (Blin et al., 2017) to identify the scytocyclamide biosynthetic gene cluster. AntiSMASH recognized 9 NRPS/PKS coding regions in the draft genome. The NRPS gene domain organization was compared to the scytocyclamide structure and neighboring candidate pathways for scytocyclamide biosynthesis were identified. Flanking genes with the same orientation to the NRPSs were included in the candidate cluster between 3,716,086-3,812,822 bp. The cluster is limited from both sides by genes with opposite orientation. Adenylation domain substrate specifity prediction was performed by combining differring AntiSMASH 4.1 and AntiSMASH 5.1.2 (Blin et al., 2019) results. The scytocyclamide biosynthetic gene cluster was visualized using Artemis (Rutherford et al., 2000) and functional annotations (Table S1) were manually refined using a combination of BLASTp and CDD database searches.

The condensation domain of NRPS module LxaC_3_ was analyzed with Natural Product Domain Seeker NaPDos (Ziemert et al., 2012) to study the role of the condensation domain in Dhb modification. The phylogenetic comparison was made with condensation domains with a similar position to Dhb in hassallidin biosynthesis (Vestola et al., 2014) and nodularin biosynthesis (Jokela et al., 2017) with the condensation domains of HasO_2_ and NdaA_1_, respectively.

## Results

### Structure of scytocyclamides

UPLC-QTOF analysis of *S. hofmannii* PCC 7110 methanol extract yielded six peaks corresponding scytocyclamide variants (Figure S1, Figure S2). Three of these (scytocyclamides A-C) have been previously characterized with spectrometric methods, including NMR. Three new less abundant variants, scytocyclamides A2, B2, and B3 appeared to be less hydroxylated (Figure 1,Table 3). The protonated masses and relative intensities for each compound are shown in Table 4. Product ion spectra (MS^E^) of protonated scytocyclamides A-C showed that the amino acid sequence could be generated from high-intensity ions in which proline is N-terminal (Figure S3, S4). Application of this fragmentation behavior to product ion spectra (MS^E^) of the new scytocyclamides A2, B2, and B3 clearly showed the amino acids lacking a hydroxyl group (Figure S3, S4). Scytocyclamides A and A2 fall in 11-residue laxaphycins and scytocyclamides B-C fall in 12-residue laxaphycins. The yields for each compound were 1 mg (A), 1 mg (A2), 3 mg (B), 0.8 mg (C), 0.4 mg (B2), and 0.4 mg (B3).

**Table 3.**
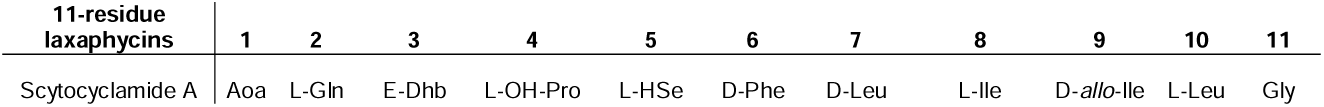
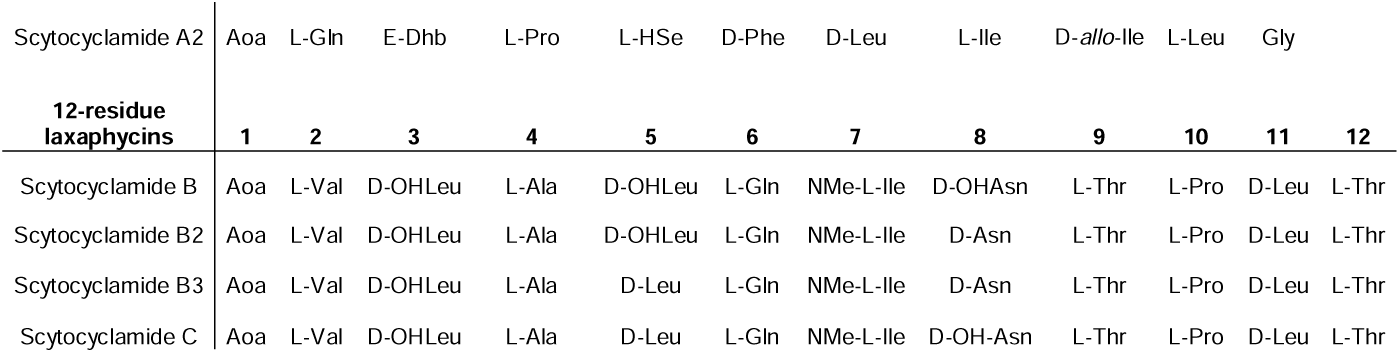
Structures of scytocyclamides from *S. hoffmannii* PCC 7110, with new variants A2, B2, and B3. Stereochemistry according to epimerase location in the biosynthetic gene cluster modules and Grewe 2005.

| 11-residue laxaphycins | 1 | 2 | 3 | 4 | 5 | 6 | 7 | 8 | 9 | 10 | 11 |
| --- | --- | --- | --- | --- | --- | --- | --- | --- | --- | --- | --- |
| Scytocyclamide A | Aoa | L-Gln | E-Dhb | L-OH-Pro | L-Hse | D-Phe | D-Leu | L-Ile | D-*allo*-Ile | L-Leu | Gly |

|  |  |  |  |  |  |  |  |  |  |  |  |  |
| --- | --- | --- | --- | --- | --- | --- | --- | --- | --- | --- | --- | --- |
| Scytocyclamide A2 | Aoa | L-Gln | E-Dhb | L-Pro | L-HSe | D-Phe | D-Leu | L-Ile | D-*allo*-Ile | L-Leu | Gly |  |
| **12-residue**<br>**laxaphycins** | **1** | **2** | **3** | **4** | **5** | **6** | **7** | **8** | **9** | **10** | **11** | **12** |
| Scytocyclamide B | Aoa | L-Val | D-OHLeu | L-Ala | D-OHLeu | L-Gln | NMe-L-Ile | D-OHAsn | L-Thr | L-Pro | D-Leu | L-Thr |
| Scytocyclamide B2 | Aoa | L-Val | D-OHLeu | L-Ala | D-OHLeu | L-Gln | NMe-L-Ile | D-Asn | L-Thr | L-Pro | D-Leu | L-Thr |
| Scytocyclamide B3 | Aoa | L-Val | D-OHLeu | L-Ala | D-Leu | L-Gln | NMe-L-Ile | D-Asn | L-Thr | L-Pro | D-Leu | L-Thr |
| Scytocyclamide C | Aoa | L-Val | D-OHLeu | L-Ala | D-Leu | L-Gln | NMe-L-Ile | D-OH-Asn | L-Thr | L-Pro | D-Leu | L-Thr |

**Table 4.** Scytocyclamides A-C from *S. hoffmannii* PCC 7110. Retention times (t_R_), experimental (Exp) mass of protonated scytocyclamides, difference (Δ) to calculated mass, chemical formula, and relative intensity (RI) of pronated scytocyclamides.

| | $t_R$ | [M+H]<sup>+</sup> | | | |
| --- | --- | --- | --- | --- | --- |
| 11-residue laxaphycins | (min) | Exp (*m/z*) | $\Delta$ (ppm) | Formula | RI (%) |
| Scytocyclamide A | 3.46 | 1223.7399 | 0,0 | C<sub>61</sub>H<sub>99</sub>N<sub>12</sub>O<sub>14</sub> | 98 |
| Scytocyclamide A2 | 3.56 | 1207.7422 | -2.3 | C<sub>61</sub>H<sub>99</sub>N<sub>12</sub>O<sub>13</sub> | 2 |
| 12-residue laxaphycins |  |  |  |  |  |
| Scytocyclamide B | 3.10 | 1367.8173 | 2.1 | C<sub>63</sub>H<sub>111</sub>N<sub>14</sub>O<sub>19</sub> | 50 |
| Scytocyclamide B2 | 3.14 | 1351.8169 | -2.0 | C<sub>63</sub>H<sub>111</sub>N<sub>14</sub>O<sub>18</sub> | 18 |
| Scytocyclamide B3 | 3.27 | 1335.8228 | -1.4 | C<sub>63</sub>H<sub>111</sub>N<sub>14</sub>O<sub>17</sub> | 10 |
| Scytocyclamide C | 3.23 | 1351.8190 | -0.4 | C<sub>63</sub>H<sub>111</sub>N<sub>14</sub>O<sub>18</sub> | 22 |

**Figure 1.**
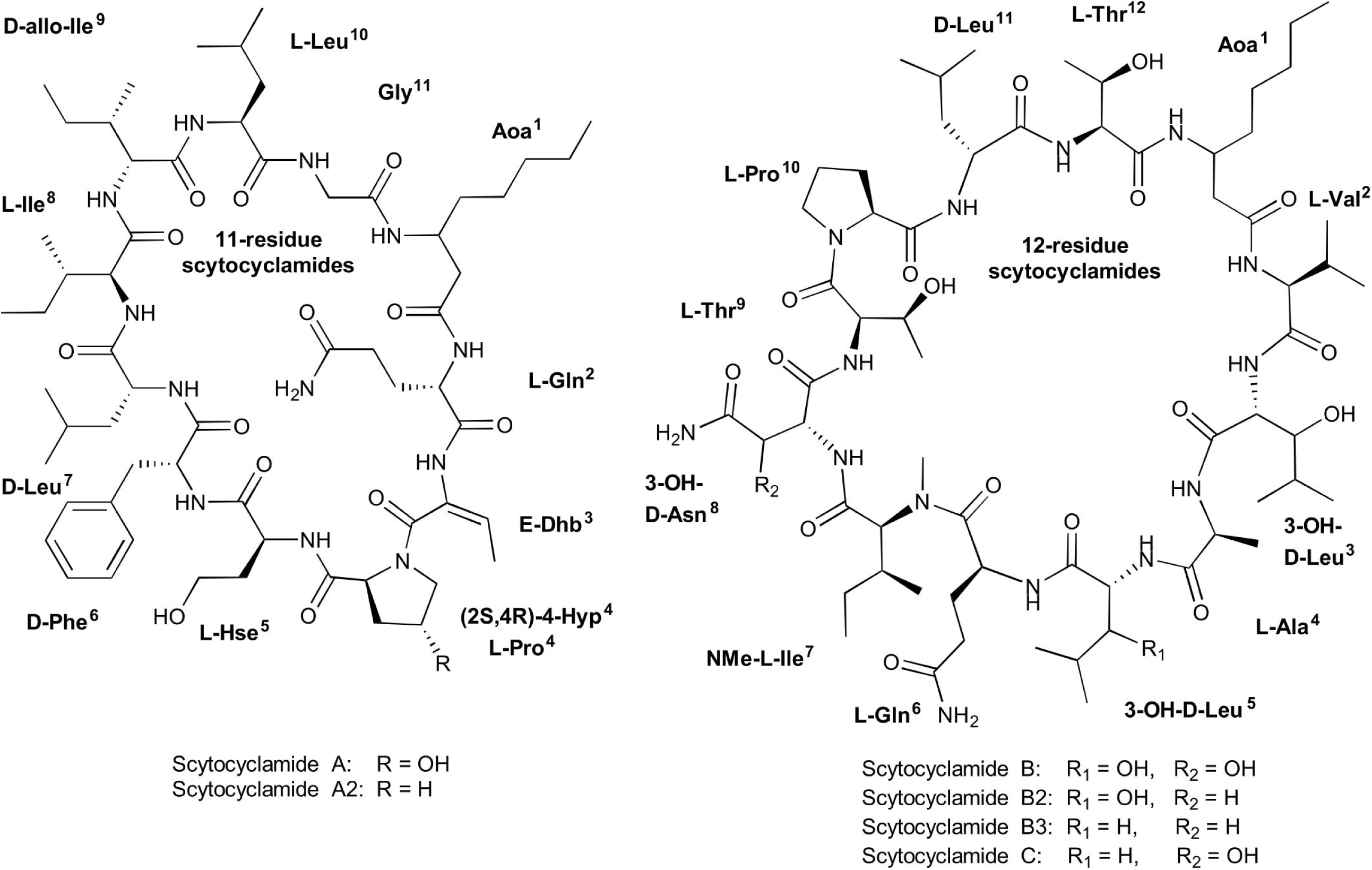
Structures of 11- and 12-residue laxaphycin variants scytocyclamides.

### Scytocyclamide biosynthetic gene cluster

Analysis of the public 12.3-Mb draft genome of *S. hofmannii* PCC 7110 identified 15 putative NRPS/PKS pathways in 9 regions recognized by AntiSMASH. Two sets of NRPSs with domain architecture matching the amino acid sequences of the two scytocyclamide types were found, separated from each other by only 5 ORFs in a 9-kb region in between them (Figure 2). Only a single pathway candidate for the initiation of the biosynthetic pathway with the fatty acid incorporation was found clustered with the NRPS genes (Figure 2). Both types of scytocyclamides contain β-aminooctanoic acid (Aoa) in their structures, and we predict that the two compounds share the initiating biosynthetic genes and pathway for the production of Aoa. The 96-kb biosynthetic gene cluster has 13 reading frames that were annotated *lxaA-H, lxbA-D*, and ORF1 (Figure 2, Table S1).

**Figure 2.**
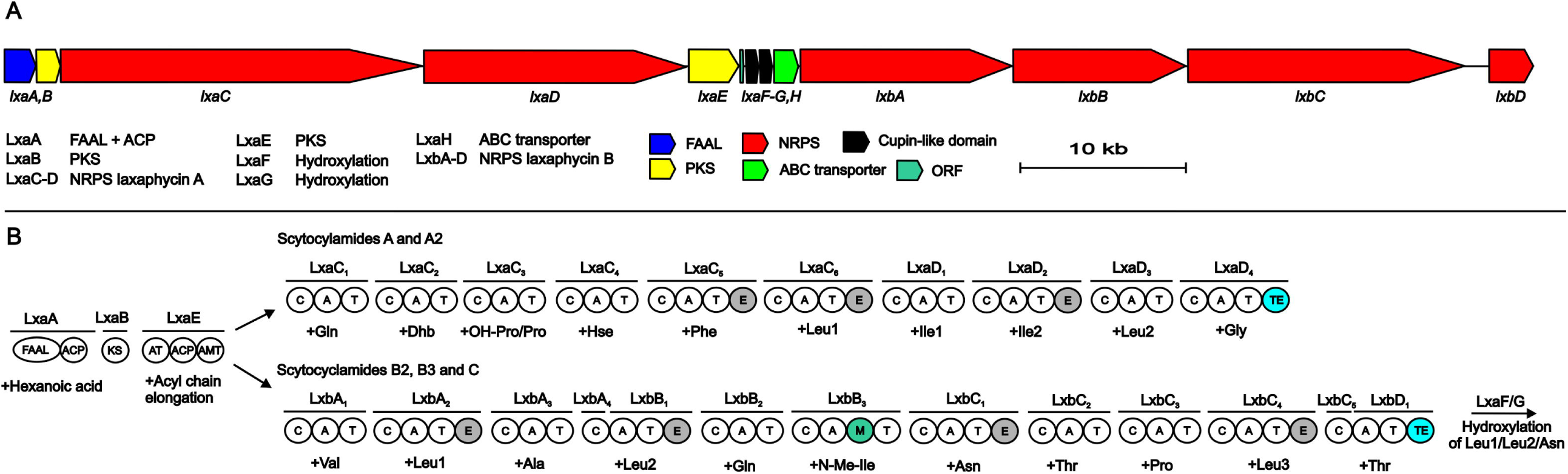
The scytocyclamide (*lxa* and *lxb*) biosynthetic gene cluster and putative biosynthetic scheme. A: Organization of predicted scytocyclamide iosynthetic genes. B: Proposed biosynthetic pathway of scytocyclamides. NRPS Non-ribosomal peptide synthetase, PKS Polyketide synthase, FAAL Fatty acyl AMP Ligase, ACP acyl carrier protein, KS ketosynthase, AT acyltransferase, AMT aminotransferase, C condensation domain, A adenylation domain, T thiolation domain, M methylation domain, TE thioesterase domain.

The predicted biosynthesis of both scytocyclamide types is initiated by the LxaA enzyme containing FAAL and ACP domains and is predicted to activate and load a hexanoic acid (Figure 2). The hexyl group is forwarded to the PKS enzymes LxaB and LxaE (Figure 2). LxaB contains a single ketosynthase (KS) domain and LxaE is composed of acyl transferase (AT), ACP, and aminotransferase (AMT) domains (Figure 2). These PKS domains elongate the hexyl chain with one acyl group to octyl chain and the aminotransferase acts on the carbonyl in the β position adding the amino group (Figure 2). We predict that β-aminooctanoic acid has two alternative branched pathways, the 11- or 12-residue scytocyclamide NRPSs (Figure 2). In 11-residue scytocyclamide synthesis the LxaC-D NRPSs and in 12-residue scytocyclamides the LxaA-D NRPS enzymes incorporate the amino acids (Figure 2). Both pathways have a terminal thioesterase (TE) that head-to-tail cyclize the compound. Each module of LxaC-D and LxbA-D enzymes bears a condensation (C), adenylation (A), and thiolation (T) domain (Figure 2). In addition, LxaC_5_, LxaC_6_, LxaD_2_ and LxbA_2_, LxbB_1_, LxbB_3_, and LxbC_4_ modules contain epimerase domains and LxbB_3_ contains a N-methylation domain (Figure 2). LxaH is an ABC-transporter characteristic to NRPS gene clusters.

The predicted adenylation domain substrate specificities of LxaC-D and LxbA-D match with the amino acids incorporated to scytocyclamides (Table S2) with some modifications. The scytocyclamide chemical structures contain 3-OHLeu, 3-OHAsn, 4-OHPro, and Dhb (Table 3). Scytocyclamide chemical variants with hydroxylations are the most abundant products produced by *S. hofmannii* PCC 7110 (Table 4). The leucine-binding pockets are identical (DAWFLGNVVK) for all four scytocyclamide leucins (position 10 in 11-residue scytocyclamides and positions 3, 5, and 11 in 12-residue scytocyclamides) with the possible exception of a gap in sequence of position 3 (---FLGNVVK) (Table S2). Cultivation of *S. hofmannii* PCC 7110 in modified growth medium containing racemic 3-OHLeu did not result in an increase of the relative amounts of hydroxylated leucine-containing laxaphycin variants (Figure S5). This could indicate that LxbA_2_ and LxbB_1_ adenylation domains incorporate Leu and not 3-OHLeu, assuming that 3-OHLeu is taken up by the cell. *S. hofmannii* PCC 7110 incorporated the non-proteinogenic amino acids (2S,4R)-MePro, (2R,4R)-MePro, (2S,4S)-MePro, (2S,4S)-OHPro, and (2S,4R)-4-OHPro in parallel cultivation experiments (data not shown). We predict that the cupin 8 family proteins LxaF-G hydroxylate the leucines and the asparagine after incorporation of the proteinogenic amino acids (Figure 2). We did not find suitable candidate enzymes for modification of hydroxyproline encoded in the BGC.

Phylogenetic analysis of the Dhb-tailoring related condensation domains LxaC_4_, HasO_2_, and NdaA_1_ with NaPDoS resulted in all of the submitted sequences having the highest identity with the modified AA clade of condensation domains (Figure S6). The modified AA clade of condensation domains have been proposed to have an active role in threonine dehydration in NRPS synthesis and our result supports this proposal

### Antimicrobial activity

Methanol extracts of *S. hofmannii* PCC 7110 inhibited the growth of *A. flavus* FBCC 2467. Disc diffusion assays were performed after purification of the scytocyclamides from the extract. Inhibition of fungal growth was observed with individual scytocyclamides as a hazy inhibition zone and synergy was observed between 11-residue and 12-residue compounds as a noticeably increased clear inhibition zone (Figure 3). Scytocyclamide amounts and inhibition zone diameters are shown in Table S3. Cross-contamination between purified scytocyclamides A-D was from <1% to 5 % and 15% for E (Figure S7).

**Figure 3.**
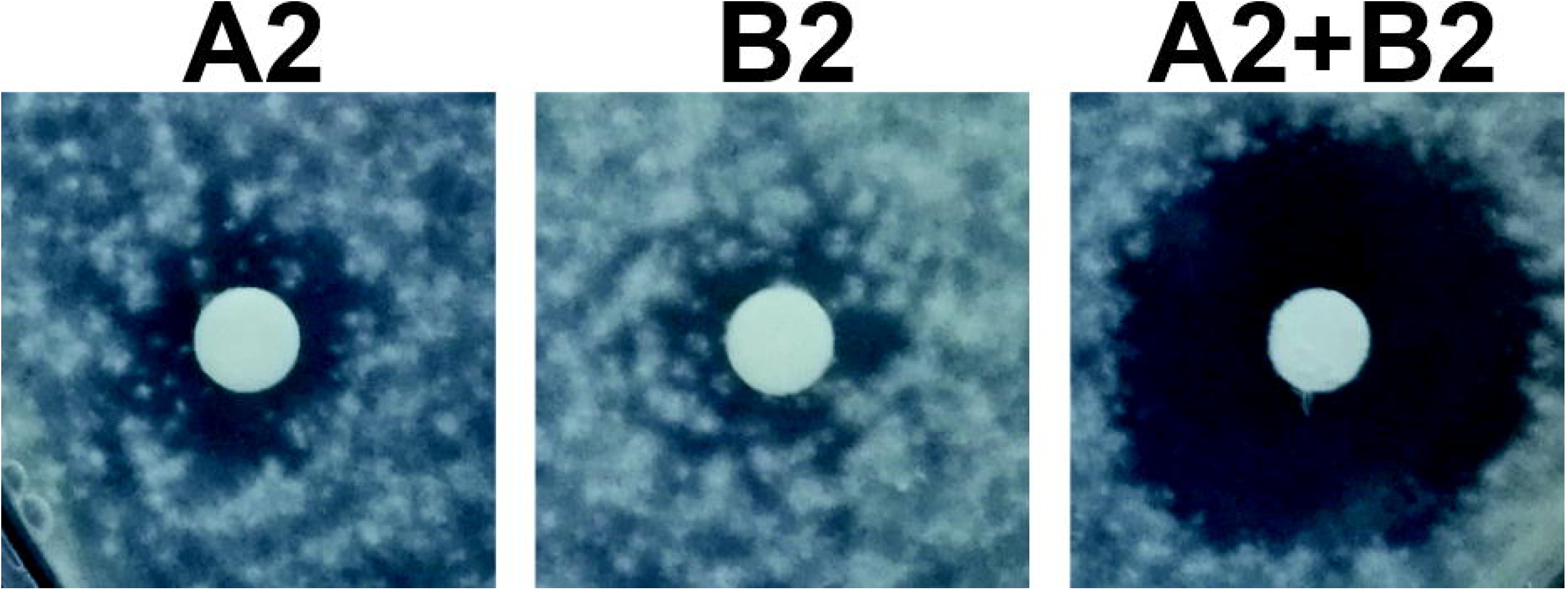
Inhibition of growth of *Aspergillus flavus* by scytocyclamides. Scytocyclamide A2 (200 *µ*g), scytocyclamide B2 (85 *µ*g), and scytocyclamides A2+B2 (100 *µ*g + 43 *µ*g). Disc diameter is 5 mm.

## Discussion

We described an unusual natural product biosynthetic gene cluster for producing structurally distinct scytocyclamides. Our analysis suggests that scytocyclamides have branched biosynthesis due to the shared loading modules LxaA-B and LxaE (Figure 2, Table S1). These shared loading modules initiate the biosynthesis with the β-amino acid Aoa, which is the only common amino acid in the peptide sequence of the two types of scytocyclamides. The biosynthesis then branches to two NRPS pathways (Figure 2). The organization of the catalytic domains in the NRPS enzymes LxaC-D matches the structure of 11-residue scytocyclamides A and A2 and NRPSs LxbA-D match the structure of 12-residue scytocyclamides B, B2, B3, and C (Figure 2), as analyzed in this study and reported earlier (Grewe, 2005). This kind of branching is exceptional to natural product biosynthetic gene clusters that are typically self-contained and act independently following the co-linearity rule of PKS/NRPS biosynthesis (Guenzi et al., 1998; Callahan et al., 2009; Baral et al., 2018). However, there are exceptions to this rule. Encoding genes are not always in a successive order (Mootz et al., 2002; Callahan et al., 2009). Modules can be skipped, as in the case of anabaenopeptin and namalide synthesis in *Nostoc* sp. CENA543, where the two compounds are produced by the same gene cluster, but a shorter product namalide is produced when three modules are skipped (Shishido et al., 2017). For example, PKS domain skipping occurs in the synthesis of leinamycin (Tang et al., 2006). Alternative starter modules have been found in the synthesis of anabaenopeptins (Rouhiainen et al., 2010) and puwainaphycins and minutissamides (Mareš et al., 2019). Gene clusters have also been shown to share enzymes in producing non-proteinogenic amino acids as in the case of anabaenopeptin and spumigin (Lima et al., 2017) and aeruginosin and spumigin, which results in the side product pseudoaeruginosin (Liu et al., 2015). Crosstalk between NRPS clusters has also been found in erythrochelin biosynthesis with two separate clusters sharing essential biosynthetic enzymes (Lazos et al., 2010). Some NRPSs incorporate multiple residues of the same amino acid iteratively, as in enterobactin synthesis (Shaw-Reid et al., 1999). Laxaphycin biosynthesis shared loading modules are now presented as a new exception to the colinearity rule of NRPS/PKS synthesis.

Twelve-residue scytocyclamides have hydroxylated Leu in positions 3 and 5 and hydroxylated Asn in position 8. However, the adenylation domain substrate specificity predictions are for proteinogenic Leu and Asn with a 100% match. We propose that the proteinogenic amino acids act as substrates for the NRPS enzymes. In the case of OHLeu and OHAsn, the modifications occur after peptide-bond formation events. We propose that the hydroxylation of these Leu and Asn residues in all laxaphycins is performed by cupin 8-like proteins of the gene cluster. The JmjC-like cupin 8 family (pfam13621) of proteins are Fe(II) or Zn(II) and α-ketoglutarate (α-KG) dependent oxygenases and act as hydroxylases and demethylases (Hewitson et al., 2002; Markolovic et al., 2016). The enzymes working as demethylases first catalyze a hydroxylation followed by fragmentation to produce a demethylated product and formaldehyde. There are examples of hydroxylation of asparagine, aspartate, histidine, lysine, arginine, and RNA in human and animal proteins (Wilkins et al., 2018). The activity of cupin 8 is specific to the amino acid position in the peptide. The location within the supercluster suggests function in the biosynthesis of the product. To our knowledge, this kind of function of cupin 8 proteins has not been previously characterized in NRPS products. The hydroxylated amino acids occur in modules with epimerase domains. This suggests that the enzymes hydroxylating the residues are specific to D-amino acids or the epimerase domains have a role in the hydroxylation. Other mechanisms have previously been found to introduce 3-hydroxylated amino acids to NRPS products (Hou et al., 2011). α-KG-dependent oxygenases hydroxylate L-arginine in viomycin (Yin and Zabriskie, 2004), L-asparagine in daptomycin-like peptide (Strieker et al., 2007), and D-glutamine in kutzneride (Strieker et al., 2009) biosyntheses. No homologs to these enzymes were found near the scytocyclamide cluster.

Dhb is enzymatically produced from threonine recognized by the adenylation domain (Challis et al., 2000). In the case of microcystin and nodularin synthesis, the dehydration has been proposed to occur due to the active role of the following condensation domain in the process (Tillett et al., 2000; Moffitt and Neilan, 2004) and bleomycin synthesis (Du et al., 2000). These microcystin and bleomycin condensation domains have been assigned to their own clade of condensation domains as “modified AA” C-domains (Ziemert et al., 2012; Bloudoff and Schmeing, 2017). When the LxaC_3_ condensation domain was analyzed by NaPDoS, it grouped with these modified AA condensation domains. The similarity of these domains with direct contact to the modified amino acid suggests that the Dhb and Dha dehydration could be indeed catalyzed by the condensation domains in these cases. For the homoserine residues, no prediction was given by AntiSMASH 5.1. However, a previous version, antiSMASH 4.1.0, did recognize the corresponding binding pocket sequence for DLKNFGSDVK as homoserine based on the Stachelhaus code. Homoserine as an amino acid in NRPS products is less common and in cyanobacteria has been previously seen in laxaphycin family peptides and nostocyclopeptide M1 (Jokela et al., 2010). However, the biosynthesis and adenylation domains for this product have not been published. Hydroxyproline has been found in other cyanobacterial natural products, such as nostoweipeptins W1-W7 and nostopeptolides L1-L4 (Liu et al., 2014). The process of incorporating the hydroxyproline or hydroxylating the prolyl residue remain unclear.

The catalytic domain organization of the scytocyclamide gene cluster matches the laxaphycin family compound structures reported earlier. The epimerizations are conserved in 11-residue laxaphycins in positions 6, 7, and 9 and in 12-residue laxaphycins in positions 3, 5, 8, and 11. The N-methylation of the amino acid in position 7 of the 12-residue laxaphycins is also conserved. Dhb^3^ is conserved in the structures of 11-residue laxaphycins. The 3-OHLeu^3^ is conserved in 12-residue laxaphycins and 3-OHLeu^5^ and OHAsn^8^ are common in 12-residue laxaphycins (Table 1). Bornancin et al. (2019) predicted that laxaphycin gene clusters should have FAAL and PKS modules to initiate biosynthesis, because the 11-residue acyclic acyclolaxaphycins have a break just before the Aoc and cyclization would be the last step of synthesis. Bornancin et al. (2015) found acyclic 11-residue laxaphycin variants with a gap between the second and third amino acid in sequence starting with the Adc. They proposed that this gap could be where the synthesis is finished and the cyclization occurs, or that the compounds they found were cleaved by environmental agents. Our results confirm the discovered acyclic 11-residue variants could be immature products of the pathway, as the linear peptide follows the biosynthetic organization we have described. With the acyclic 12-residue variants, the gap in the sequence occurs within a predicted NRPS gene and the proposed mechanism of other agents or enzymes in the environment cleaving the products would seem more reasonable.

Cyanobacteria are abundant primary producers in aquatic environments and are targeted to grazing by higher organisms, such as sea hares (Cruz-Rivera and Paul, 2007). Cyanobacteria produce a wide range of bioactive natural products (Dittmann et al., 2015; Demay et al., 2019) that seem to be produced to deter the grazing fauna in the environment (Leão et al., 2012; Mazard et al., 2016). Potential competitors to cyanobacteria are also other microbes such as chytrids, which are fungi parasitic to cyanobacteria (Agha et al., 2018). Some cyanobacterial natural products have reached clinical trials and are approved as cancer drugs (Luesch et al., 2001; Deng et al., 2013). Cyclic lipopeptides are common among the cyanobacterial natural products and typically contain a single fatty acid as in laxaphycins (Galica et al., 2017) that confers membrane-disruptive properties (Humisto et al., 2019). Laxaphycin family peptides have been shown to be toxic to or inhibit the growth of multiple organisms and cell lines (Gerwick et al., 1989; Frankmölle et al., 1992b; Bonnard et al., 1997; MacMillan et al., 2002; Bonnard et al., 2007; Maru et al., 2010; Luo et al., 2014; Luo et al., 2015; Dussault et al., 2016; Cai et al., 2018; Bornancin et al., 2019). We observed antifungal activity of scytocyclamides towards *A. flavus* (Figure3, Table S3). In an earlier report by Grewe (2005), no activity against *C. albicans* was detected for scytocyclamides A, B, and C, which was also observed in this study. Synergistic antifungal activity between 11- and 12-residue laxaphycins has been previously reported (Frankmölle et al., 1992b; MacMillan et al., 2002). The same synergistic activity was observed between 11- and 12-residue scytocyclamides (Figure 3, Table S3). According to previous studies and our results, the 12-residue laxaphycins are typically more potent on their own than 11-residue laxaphycins. Our previous study on *S. hofmannii* PCC 7110 failed to identify the antifungal agent in the extract, when purified fractions lacked activity. We now conclude that the antifungal activity was most probably caused by scytocyclamides, but the purified fractions had insufficient amounts of material to produce the inhibitory effect without a synergistic partner (Shishido et al., 2015).

It is probable that the other type of laxaphycins originally existed without a synergistic partner peptide in the cells, as many laxaphycins have antimicrobial activity by themselves. Through recombination events, a synergistically acting peptide has emerged to enhance the activity of the original peptide. One possibility is that the two peptides had individual gene clusters, but the initiating enzymes have been subject to an elimination event when two distinct starter enzymes were no longer necessary. It is clear that the synergistic bioactivity and shared biosynthesis of laxaphycins go together. Similar colocalization with coregulation of distinct synergistic biosynthetic gene clusters has been previously observed in the streptomycetal antibiotics griseoviridin and viridogrisein (Xie et al., 2012). The mechanism behind the synergistic action is usually two different compounds acting on two different targets, thus combining their activity (Caesar and Cech, 2019). It is possible that one compound makes the target cell vulnerable to the other, such as via damage to the cell wall. The colocalization of genes and shared biosynthesis suggest simultaneous regulation and expression of the synergistic products to act on a single cellular target through different mechanisms.

## Author Contributions

LMPH, KS, JJ, MW, and DPF designed the study.

AJ, LMPH, and MW performed the experiments.

LMPH, JJ, and DPF analyzed and interpreted the data.

LMPH, DPF, JJ, and KS wrote the manuscript, which was corrected, revised, and approved by all authors.

## Acknowledgments

The authors thank Lyudmila Saari for maintaining the cyanobacterial strain retrieved from The Pasteur Culture Collection of Cyanobacteria.

