## Supplementary material for "Shared PKS modules in biosynthesis of synergistic laxaphycins"

Table S1. Predicted functions of the ORFs in the *lxa* cluster.

| **Protein** | **Accession number** | **Length (AA)** | **Function** | **Similarity (Protein, origin)** | **Pairwise identity** | **Accession number** |
| --- | --- | --- | --- | --- | --- | --- |
| **LxaA** | WP_017742670.1 | 715 | FAAL+ACP | PuwI *Symplocastrum muelleri* NIVA-CYA 644 | 59.72 | AXN93619.1 |
| **LxaB** | WP_017742671.1 | 474 | KS | PuwB *Anabaena minutissima* UTEX B 1613 | 71.18 | AXN93597.1 |
| **LxaC** | WA1_RS15505^a^ | 7398 | NRPS | non-ribosomal peptide synthetase *Tumebacillus avium* | 45.15 | WP_087458112.1 |
| **LxaD** | WP_017742673.1 | 5058 | NRPS | non-ribosomal peptide synthetase *Nostoc flagelliforme* | 66.91 | WP_100898072.1 |
| **LxaE** | WP_066612891.1 | 1161 | AT+ACP+AMT | PuwE [Symplocastrum muelleri NIVA-CYA 644] | 54.71 | AXN93622.1 |
| **ORF1** | WP_017742675.1 | 58 | Hypotethical protein | hypothetical protein *Scytonema* sp. UIC 10036 | 52.78 | WP_155750574.1 |
| **LxaF** | WP_017742676.1 | 296 | Cupin 8 | cupin-like domain-containing protein *Tolypothrix bouteillei* | 74.23 | WP_050045604.1 |
| **LxaG** | WP_017742677.1 | 305 | Cupin 8 | cupin-like domain-containing protein *Tolypothrix bouteillei* | 78.21 | WP_050045604.1 |
| **LxaH** | WP_017742678.1 | 595 | ABC transporter | ABC transporter ATP-binding protein/permease *Tolypothrix bouteillei* | 93.45 | WP_038095421.1 |
| **LxbA** | WP_148662958.1 | 4457 | NRPS | non-ribosomal peptide synthetase *Nostoc flagelliforme* | 64.50 | WP_100898072.1 |
| **LxbB** | WP_148662959.1 | 3434 | NRPS | non-ribosomal peptide synthetase *Nodularia* sp. NIES-3585 | 78.88 | WP_089089654.1 |
| **LxbC** | WP_066612897.1 | 5742 | NRPS | non-ribosomal peptide synthetase *Tolypothrix bouteillei* | 80.23 | WP_050045607.1 |
| **LxbD** | WP_066612900.1 | 839 | NRPS | non-ribosomal peptide synthetase *Tolypothrix bouteillei* | 89.92 | WP_063779483.1 |

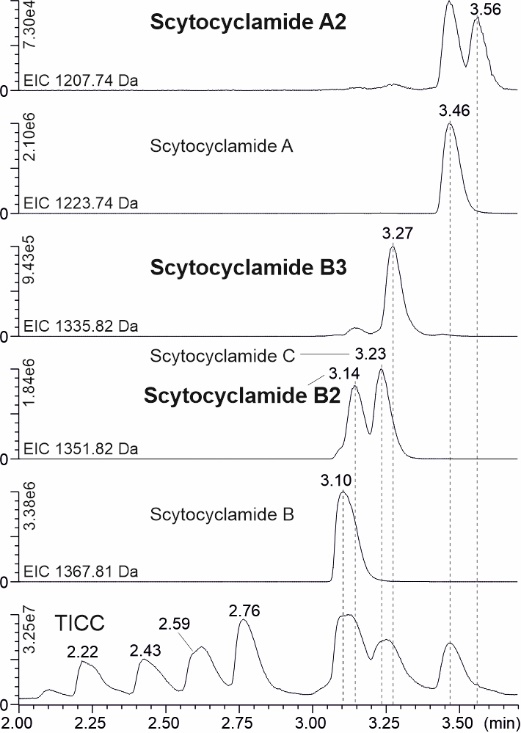

Figure S1. Extracted ion chromatograms (EIC) of scytocyclamides and total ion current chromatogram (TICC). Names of new variants in bold.

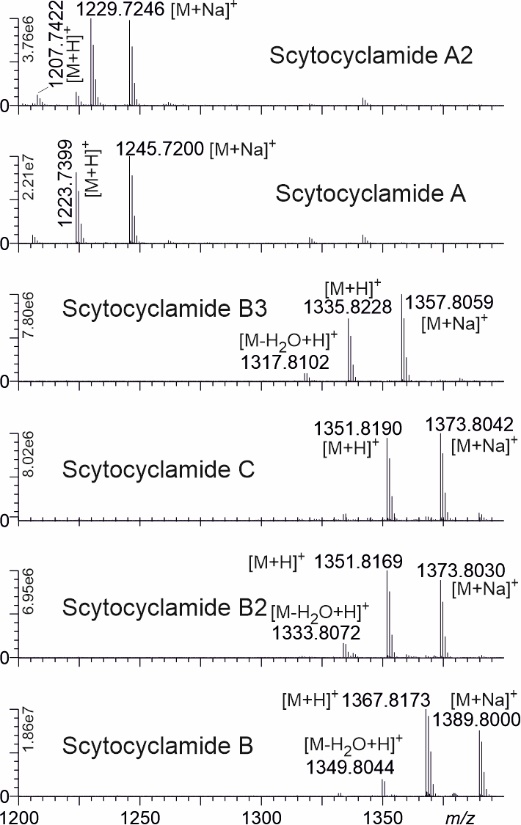

Figure S2. Mass spectra from scytocyclamide peaks with [M+H]^+^ and [M+Na]^+^ ion masses.

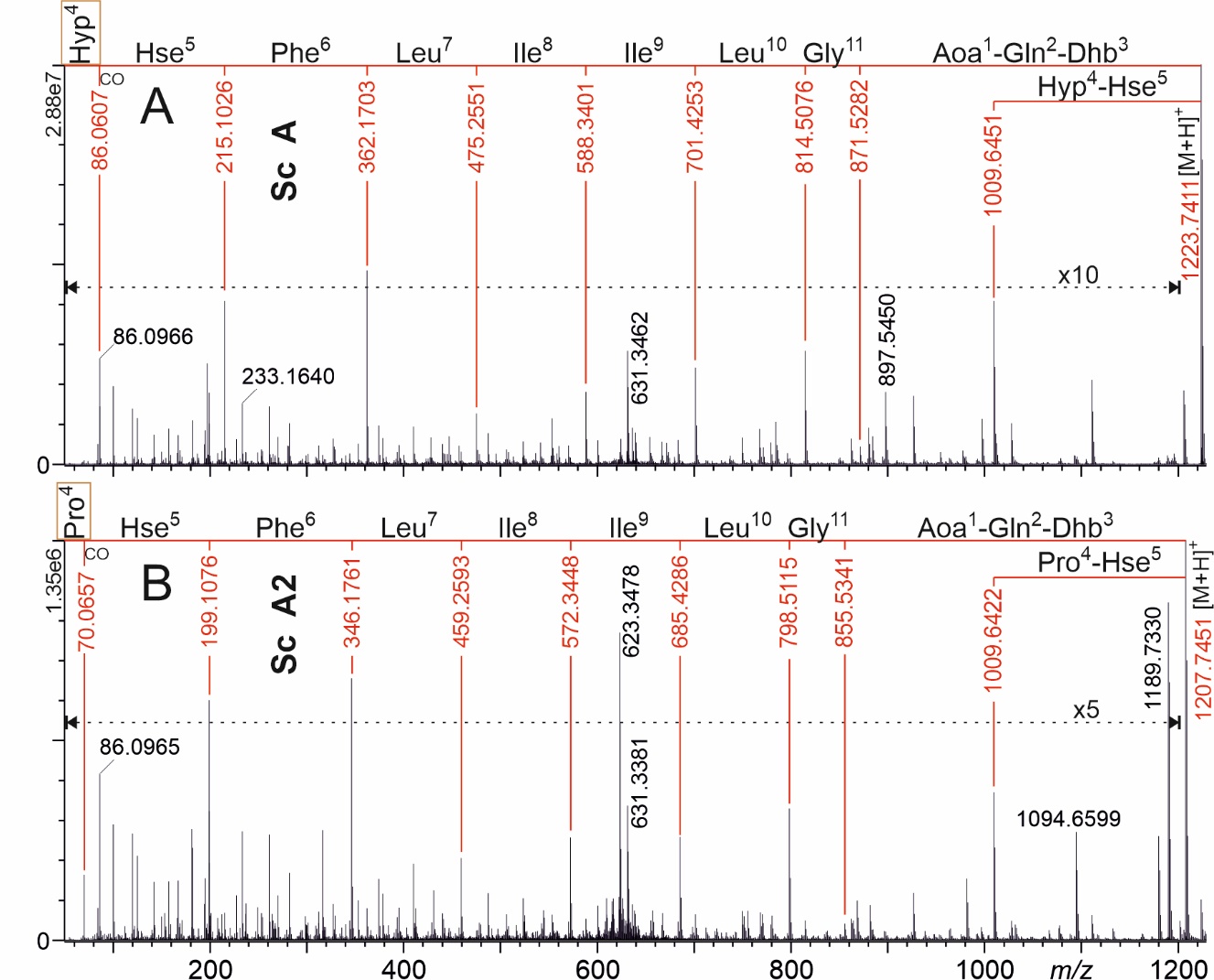

Figure S3. UPLC-QTOF product ion mass spectra of protonated 11-residue scytocyclamides (Sc) A (A) and A2 (B). The most complete product ion series showing the amino acid sequences are marked with red numbers and lines. Hyp = OHPro, Hse = Homoserine.

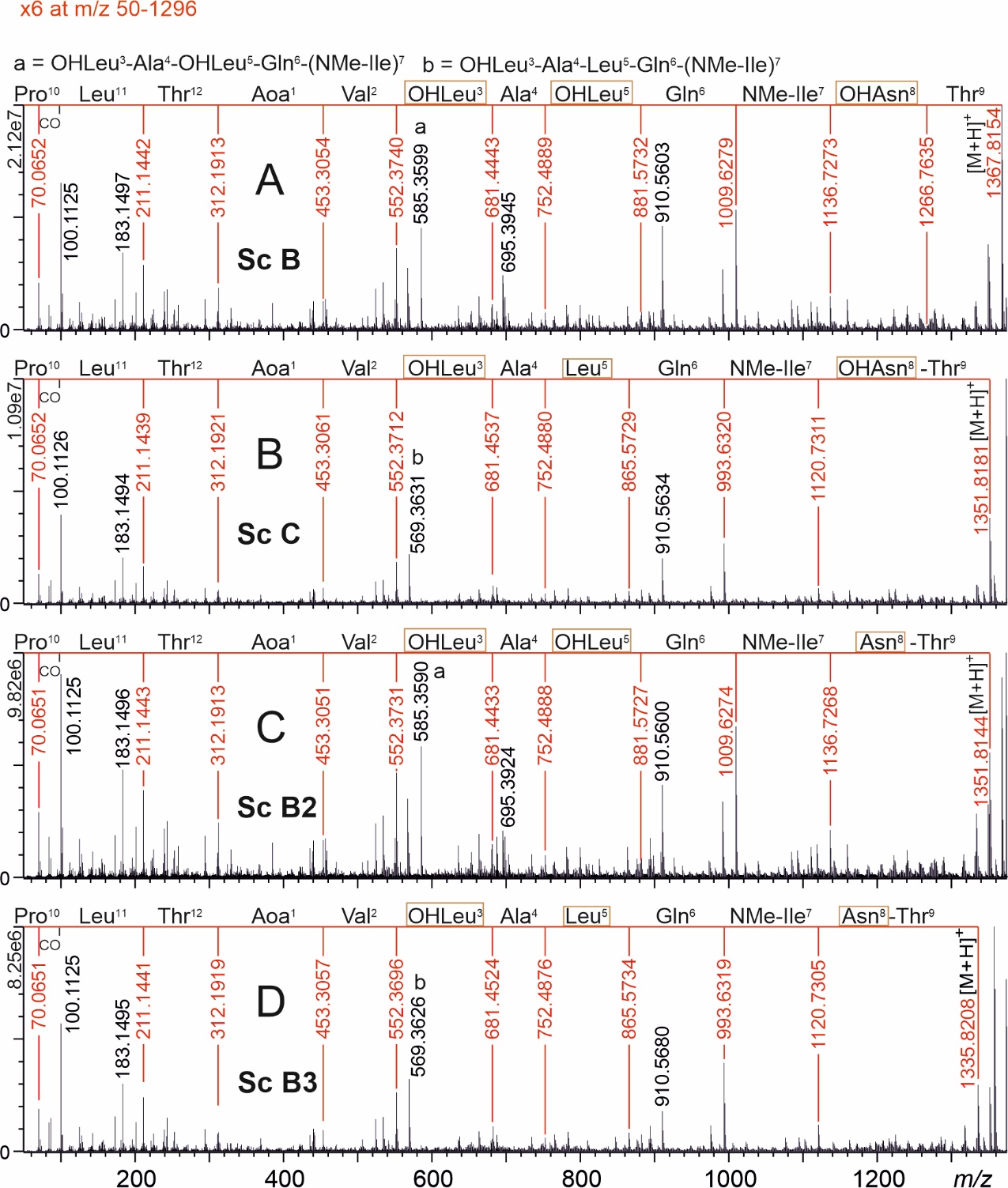

Figure S4. UPLC-QTOF product ion mass spectra of protonated 11-residue scytocyclamides (Sc) B (A), C (B), B2 (C), and B3 (D). The most complete product ion series showing the amino acid sequences are marked with red numbers and lines.

Table S2. Predicted and activated substrates of adenylation domains in scytocyclamids, with the binding pocket amino acid residues as identified with NRPSpredictor2 in AntiSMASH 5.1.2

| **AA residue** | **Protein** | **Predicted** | **Activated** | **A-domain residues** | **Stachelhaus code % match** |
| --- | --- | --- | --- | --- | --- |
|  | **11-residue scytocyclamides** | | | | |
| 2 | LxaC_1_ | Gln | Gln | DAWQFGLIDK | 100 |
| 3 | LxaC_2_ | Thr | Thr | DFWNIGMVHK | 100 |
| 4 | LxaC_3_ | Pro | Hyp/Pro | DahFIAHVVK | 80 |
| 5 | LxaC_4_ | Hse* | HSe | DlknFGSdvK | * |
| 6 | LxaC_5_ | Phe | Phe | DAWtIAAVCK | 90 |
| 7 | LxaC_6_ | Leu | Leu | DAWFLGNVVK | 100 |
| 8 | LxaD_1_ | Ile | Ile | DAFFLGVTFK | 100 |
| 9 | LxaD_2_ | Ile | Ile | DAFFLGVTFK | 100 |
| 10 | LxaD_3_ | Leu | Leu | DAWFLGNVVK | 100 |
| 11 | LxaD_4_ | Gly | Gly | DILQLGLIWK | 100 |
|  | **12-residue scytocyclamides** | | | | |
| 2 | LxbA_1_ | Val | Val | DAfWLGGTFK | 90 |
| 3 | LxbA_2_ | Leu | Leu | DAWFLGNVVK | 100 |
| 4 | LxbA_3_ | Ala | Ala | DLFNNALTYK | 100 |
| 5 | LxbB_1_ | Leu | Leu | ---FLGNVVK | 70 |
| 6 | LxbB_2_ | Gln | Gln | DAWQFGLIDK | 100 |
| 7 | LxbB_3_ | Ile | Ile | DAFFLGVTFK | 100 |
| 8 | LxbC_1_ | Asn | Asn | DLTKIGEVGK | 100 |
| 9 | LxbC_2_ | Thr | Thr | DFWNIGMVHK | 100 |
| 10 | LxbC_3_ | Pro | Pro | DVQFmAHVVK | 90 |
| 11 | LxbC_4_ | Leu | Leu | DAWFLGNVVK | 100 |
| 12 | LxbD_1_ | Thr | Thr | DFWNIGMVHK | 100 |

*Prediction by AntiSMASH 4.1.0 based on stachelhaus code, no % match given by the program.

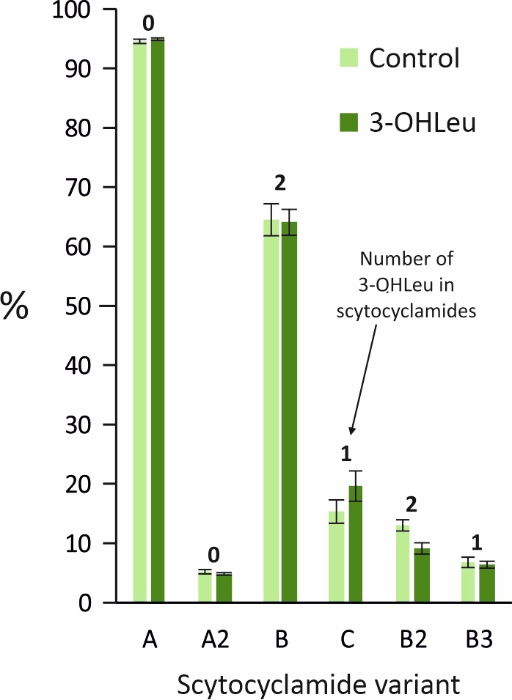

Figure S5. Relative (%) intensities of 11- (A, A2) and 12- (B, C, B2, B3) residue scytocyclamides (sum of single and doubly charged protonated and sodiated ions) from cultivations with (3-OHLeu) and without (Control) added racemic 3-hydroxyleucines in growth medium. Slight changes in the intensities can be seen, but the changes do not favor variants with more 3-OHLeu. Relative scytocyclamide intensities in Table 4 differ from these control values due to different growth conditions in the mass cultivations.

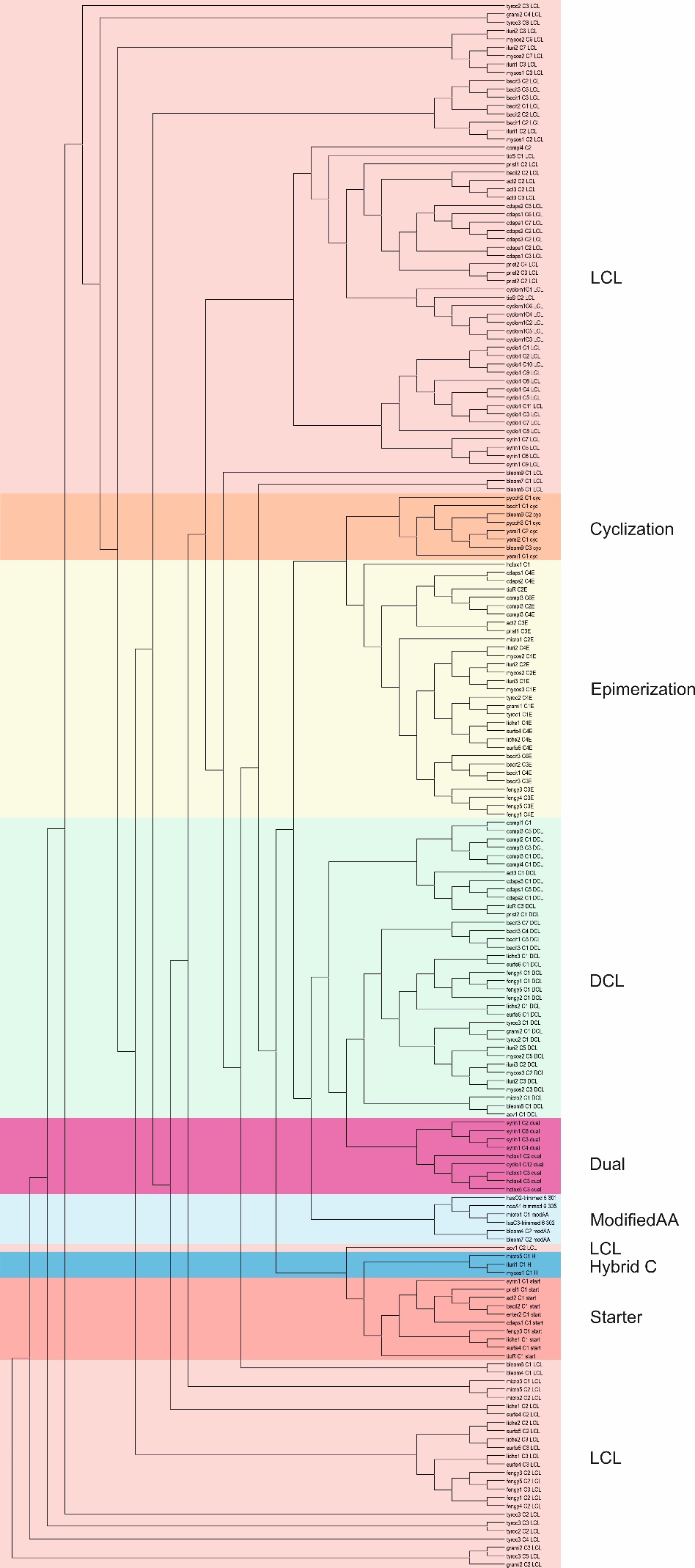

Figure S6. Phylogeny-based C-domain classification, with LxaC_4_, HasO_2_, and NdaA_1_ C-domains clustered in the modified AA clade. The phylogenetic tree was produced with NaPDoS (Ziemert et al., 2012). LCL - catalyzes a peptide bond between two L-amino acids, Cyclization - catalyzes peptide bond formation and cyclization, Epimerization - changes the chirality of the last amino acid, DCL - adds an L-amino acid to the peptide ending with a D-amino acid, Dual - catalyzes both epimerization and condensation reactions, ModifiedAA - involved in the modification of the incorporated amino acid, Hybrid C -involved in the condensation of an amino acid to an aminated polyketide resulting in a hybrid PKS/NRPS product, Starter - acylates the first amino acid with a β-hydroxy-carboxylic acid.

Table S3. Antifungal activities of scytocyclamides.

| **Compound(s)** | **Amount (µg)** | **Inhibition zone (mm)**  ***Aspergillus flavus*** |
| --- | --- | --- |
| **11-residue laxaphycins** |  |  |
| Scytocyclamide A | 200 | 10 |
| Scytocyclamide A2 | 200 | 7 |
| **12-residue laxaphycins** |  |  |
| Scytocyclamide B | 600 | 23 |
| Scytocyclamide C | 160 | 22 |
| Scytocyclamide B2 | 85 | 10 |
| Scytocyclamide B3 | 85 | 20 |
| **Synergism** |  |  |
| Scytocyclamide A+B | 100 + 300 | 36 |
| Scytocyclamide A+C | 100 + 80 | 33 |
| Scytocyclamide A2+B2 | 100 + 43 | 24 |
| Scytocyclamide A2+B3 | 100 + 43 | 25 |
| Scytocyclamide B+C | 300 + 80 | 23 |

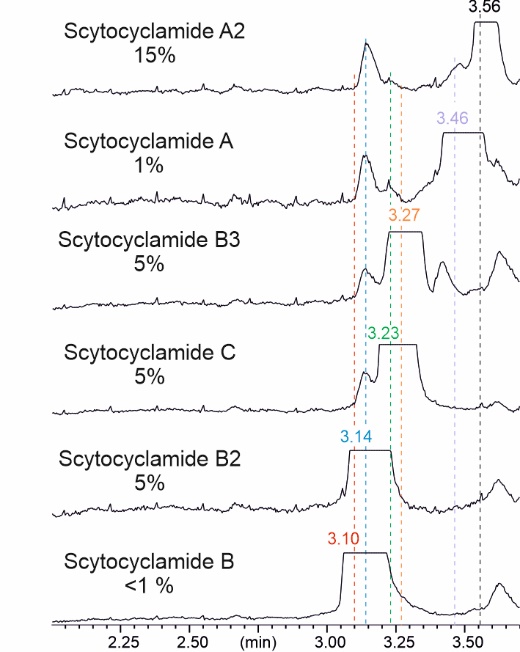

Figure S7. Total ion current chromatograms of cross-contamination between purified scytocyclamides.
